# Texture profile analysis and rheology of plant-based and animal meat

**DOI:** 10.1101/2024.10.11.617917

**Authors:** Reese A. Dunne, Ethan C. Darwin, Valerie A. Perez Medina, Marc E. Levenston, Skyler R. St. Pierre, Ellen Kuhl

## Abstract

Plant-based meat can help combat climate change and health risks associated with high meat consumption. To create adequate mimics of animal meats, plant-based meats must match in mouthfeel, taste, and texture. The gold standard to characterize the texture of meat is the double compression test, but this test suffers from a lack of standardization and reporting inconsistencies. Here we characterize the texture of five plant-based and three animal meats using texture profile analysis and rheology, and report ten mechanical features associated with each product’s elasticity, viscosity, and loss of integrity. Our findings suggest that, of all ten features, the stiffness, storage, and loss moduli are the most meaningful and consistent parameter to report, while other parameters suffer from a lack of interpretability and inconsistent definitions. We find that the sample stiffness varies by an order of magnitude, from 418.9±41.7kPa for plant-based turkey to 56.7±14.1kPa for tofu. Similarly, the storage and loss moduli vary from 50.4±4.1kPa and 25.3±3.0kPa for plant-based turkey to 5.7±0.5kPa and 1.3±0.1kPa for tofu. All three animal products, animal turkey, sausage, and hotdog, consistently rank in between these two extremes. Our results suggest that–with the right ingredients, additives, and formulation–modern food fabrication techniques can create plant-based meats that successfully replicate the full viscoelastic texture spectrum of processed animal meat.

## 1. Introduction

The global population is exponentially increasing, projected to reach nearly 10 billion by the year 2050 [12]. Consequently, the demand for food, particularly protein sources, has upsurged to meet the nutritional needs of this rapidly growing population. Since the early 1960s, global meat production has exceeded 350 million tonnes, while per capita meat consumption has approximately doubled during the same period [26]. The drastically high rates of meat production and consumption worldwide are only continuing to speed up and pose serious environmental and health risks to the planet and its inhabitants.

The ever-growing livestock industry, which employs a staggering 1.3 billion people globally [8], is responsible for 18% of global greenhouse gas emissions and has increased emission rates by 51% during the period from 1961 to 2010 [4, 8]. Moreover, pasture expansion for the beef industry alone accounts for 41% of global deforestation [27]. Global greenhouse gas emissions from livestock, fertilization, and deforestation play major roles in climate change, biodiversity loss, water pollution, and global warming [2, 8, 28]. In addition to environmental consequences, studies show that meat consumption, particularly of red meat, is linked to cardiovascular disease, type II diabetes, cancer, and other adverse health complications [11]. Thus, it is crucial to explore alternative protein sources and animal meat substitutes to improve the overall health and well-being of the planet and global population.

The demand for plant-based alternatives to animal meat has risen in recent decades, not only due to health and environmen-tal concerns, but also because of ethical issues regarding animal welfare. Over 80 million animals are confined and slaughtered annually to produce the 350 million tonnes of animal meat for human consumption [8]. More people worldwide are adopting vegetarian, semi-vegetarian, or flexitarian diets as our awareness of their health and environmental benefits grows [16]. Despite studies showing that the majority of meat consumers agree with the ethical, environmental, and health viewpoints of vegetarianism and veganism [3], most consumers still prefer animal meats and are reluctant to switch to a vegetarian or vegan lifestyle [13, 20].

While taste and price are key concerns for consumers choosing between plant-based and animal-based meats [8], the main characteristic of plant-based meats that drives consumers away is the difference in texture and sensory appeal [10, 13, 36]. Plant-based meat producers have struggled to develop meat alternatives that mimic the sensory experience of animal meat [30]. Animal meats are anisotropic materials consisting of muscle fibers, connective tissue, fat, and water [17, 34], all inter-linked in a complex internal structure that is challenging for non-fibrous pea- and soy-based meat alternatives to replicate [15, 31]. Ultimately, producers need to develop meats that share the same, or similar, mechanical characteristics to persuade consumers to switch to plant-based meat substitutes [32].

Texture profile analysis is a widely known method used in food science to measure the mechanical properties of various meats and other foods [6, 24, 29]. It consists of a double compression test that consists of two cycles of loading and unloading to simulate the act of chewing. The force data from the double compression test can be used to extract an assortment of textural parameters–stiffness, hardness, cohesiveness, springiness, resilience, chewiness–that provide insight into the sensory experience of the food [24]. In contrast to our previous mechanical tests of tension, compression, and shear on tofurky, animal, and plant-based chicken at a *low strain rate* to minimize viscoelastic effects [33], the texture profile analysis is performed at a *high strain rate* to mimic the experience of chewing [7, 14]. The parameters from texture profile analyses are well correlated to sensory tests [14, 23]. However, the sample dimensions and loading rates of the texture profile analysis vary across different studies. This lack of standardization results in inconsistent definitions of the reported parameters, making cross-study comparisons difficult [14, 23, 32]. Nevertheless, performing texture profile analyses on plant-based and animal meat can highlight their differences in mechanical and textural properties, and provide consumers and food scientists with quantifiable metrics for the sensory perception of both types of meat.

Rheology, including amplitude and frequency sweep tests, can be performed to characterize the viscoelastic properties and deformation behavior of plant-based and animal meats [24]. Process parameters, for example from mixing and extrusion, can help manipulate and fine-tune the rheological properties of the final product [24]. Thus, rheological testing can provide manufacturers with valuable information on how to process plant-based meats to achieve similar properties as animal meat. Combining rheology with texture profile analysis provides valuable information on the elastic and viscous components of meat, which collectively contribute to our textural and sensoral perception [24].

Here we use texture profile analysis and rheology to quantify the elastic, viscous, and textural properties of eight commercially available plant-based and animal meats, which we have previously characterized in tension, compression, and shear and in a food texture survey [31]. We explore to which extent the parameters from the texture profile analysis and rheological tests are correlated to classical mechanical testing and to our sensory perception of texture.

## 2. Methods

### 2.1. Sample preparation

We test *n* = 5 plant-based meats, Plant-Based Signature Stadium Hot Dog (Field Roast, Seattle, WA), Vegan Frankfurter Sausage (Field Roast, Seattle, WA), Organic Firm Tofu (365 by Whole Foods, Austin, TX), Organic Tofu Extra Firm (House Foods, Garden Grove, CA), and Ham-Style Roast Tofurky (Tofurky, Hood River, OR). For comparison, we test *n* = 3 animal meats, Classic Uncured Wieners (Oscar Mayer, Kraft Heinz Co, Chicago, IL), Turkey Polska Kielbasa Sausage (Hillshire Farm, New London, WI), and Spam Oven-Roasted Turkey (Spam, Hormel Foods Co, Austin, MN). We select these products motivated by our previous studies to directly compare mechanical testing and sensory perception [31] to the texture profile analysis and rheological tests commonly used in food science research [32]. All meats of our study are comminuted products [18]. Unlike whole meats, we do not expect them to have natural, large length-scale anisotropic properties [19]. Table 3 summarizes the brand name, manufacturer, and list of ingredients of all eight products. For all eight meats, we use a biopsy punch to extract cylindrical samples of 8 mm diameter and 10 mm height from the center of each product. We confirm the sample diameter at three locations along the height and store all samples at room temperature at 25^°^ C for a maximum of 30 minutes until testing. Figure 1 shows the sample preparation process.

**Figure 1:**
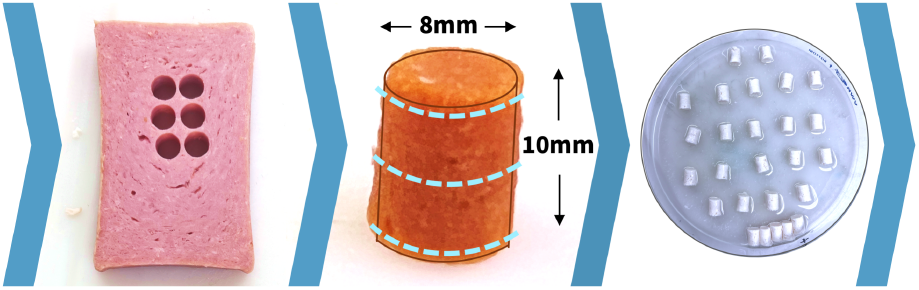
Sample preparation. We test *n* = 5 plant-based and *n* = 3 animal meats in compression and shear. To prepare the samples, we use a biopsy punch and extract cylindrical samples with 8 mm diameter and 10 mm height from the center of each product, left. We confirm the sample diameter at three locations along the height, middle, and store all samples at room temperature until testing, right.

### 2.2. Sample testing

For each of the eight products, we prepare *n* = 13 samples for uniaxial double compression testing, *n* = 1 sample for initial amplitude sweep shear testing, and *n* = 5 samples for frequency sweep shear testing. We perform all compression and shear tests using an AR-2000ex torsional rheometer (TA Instruments, New Castle, DE). Figure 2 displays our compression and shear test setup for all eight meats.

**Figure 2:**
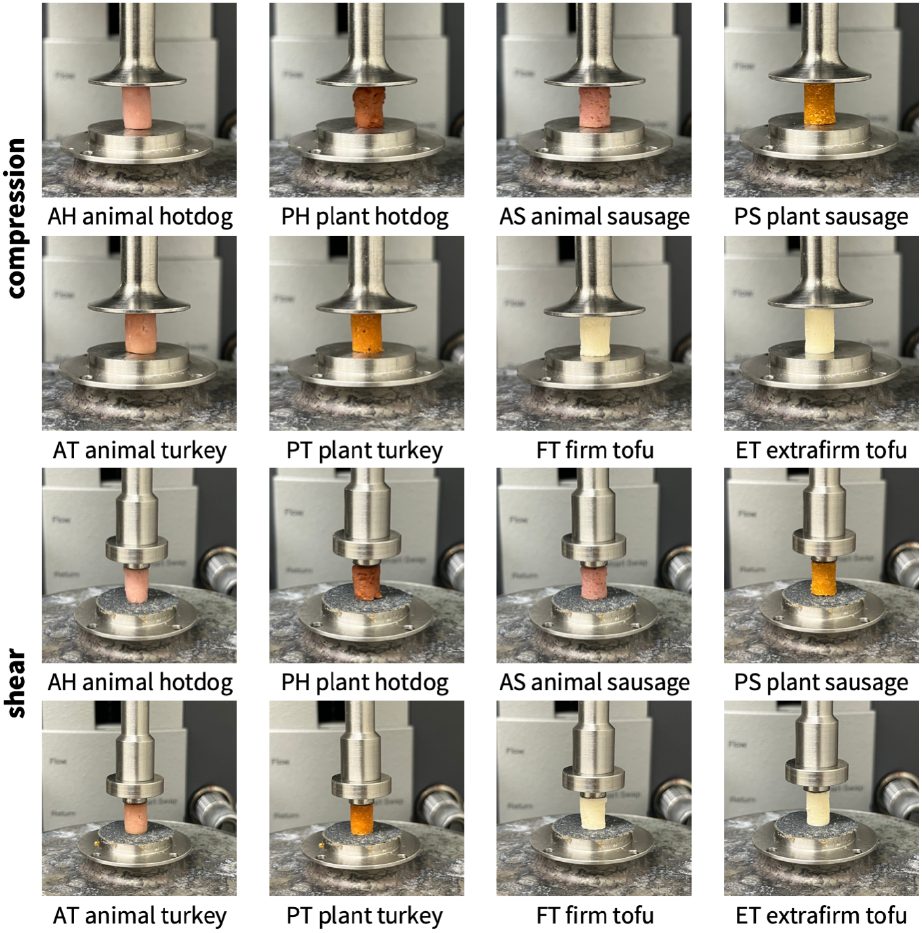
Sample testing. For all eight meats, we use *n* = 13 samples for double compression testing, *n* = 1 sample for initial amplitude sweep testing, and *n* = 5 samples for frequency sweep testing with an AR-2000ex torsional rheometer. For the double compression tests, top, we mount the samples between a 25 mm diameter base plate and a 25 mm parallel plate and compress them twice to half of their initial height. For the shear tests, bottom, we mount the samples between a 20 mm diameter base plate and a 8 mm parallel plate, both sandpaper-covered to avoid slippage, and perform frequency sweep tests.

#### 2.2.1 Compression testing for texture profile analysis

For the double compression tests, we mount the samples between a 25 mm diameter base plate and a 25 mm parallel plate and compress them to *ε* = −50% strain at the rheometer’s maximum strain rate of 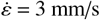, which translates to 30%/s based on the sample height. After reaching the peak strain of *ε* = −50%, we unloaded the samples at the same rate of 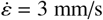, and repeated this loading and unloading process a second time. We record all compression data at a rate of 244 points/s using the fast data logger utility of the TRIOS software.

#### 2.2.2. Shear testing for rheological analysis

For the shear tests, we mount the samples between a 20 mm diameter base plate and a 8 mm parallel plate, both sandpaper-covered to avoid slippage, and compress them to *ε* = −10% strain. We perform an initial *amplitude sweep test*, oscillating at 0.5 Hz with strain amplitudes ranging from *γ* = 0.01% to *γ* = 60%. From the recordings, we determine the strain amplitude within the linear viscoelastic regime to use for the subsequent frequency sweep tests. We then perform five *frequency sweep tests* from 0.1 rad/s to 100 rad/s at approximately 4.71% strain for all meats. From the frequency sweep test, we average the storage modulus *G*′, loss modulus *G*′′, complex shear modulus *G*^*^, and phase angle *δ* between the frequency range from 0.22 to 4.64 rad/s, the plateau region on a log-log scale for all meats.

### 2.3. Texture profile analysis

The double compression tests serve as the basis for the texture profile analysis. Figure 3 illustrates the load profile of the double compression test, which consists of two consecutive cycles of 50% compression. We perform *n* = 13 double compression tests for each of the eight meats and average all 13 datasets, which results in characteristic force vs. time curves similar to the curves in Figure 3. We threshold the force values at 0.07 N minimum, which corresponds to the noise level in the raw data above which the upper rheometer plate is sufficiently in contact with the sample. According to standard definitions, we denote the peak forces of the first and second loading cycles as *F*_1_ and *F*_2_, the associated loading times as *t*_1_ and *t*_2_, the areas under their loading paths as *A*_1_ and *A*_3_, and the areas under their unloading paths as *A*_3_ and *A*_4_ [9, 1]. We convert the curves into stress vs. strain curves, where the stress *σ* = *F*/*A* is the recorded force *F* divided by the specimen cross section area *A* = *π r*^2^ = −50.3 mm^2^ with *r* = 4 mm, and the strain 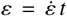 is the applied strain rate multiplied by the time *t* [32]. From the peak force of the first loading cycle *F*_1_, we calculate the peak stress *σ*_1_ = *F*_1_/*A* at peak strain *ε* = −50%. From these characteristic values, we extract six texture profile analysis parameters [22]. The *stiffness E* = *σ*/*ε* refers to the slope of stress-strain curve during first compression, when the specimen is compressed to half of its height. The *hardness F*_1_ is associated with the peak force during this first compression cycle. The *cohesiveness* (*A*_3_ + *A*_4_)/(*A*_1_ + *A*_2_) characterizes the material integrity during the second loading and unloading cycle, compared to the first cycle, where a value of one relates to a perfectly intact material, whereas a value of zero relates to complete disintegration. The *springiness t*_2_/*t*_1_ is associated with recovery and viscosity and describes the speed by which the material springs back to its original state after the second cycle compared to the first cycle. The *resilience A*_2_/*A*_1_ is a measure of how well a sample recovers during first unloading path relative to first loading, where a value of one relates to perfect elasticity whereas a value larger than one indicates plasticity. The *chewiness F*_1_ (*A*_3_ + *A*_4_)/(*A*_1_ + *A*_2_) *t*_2_/*t*_1_, the product of hardness, cohesiveness, and springiness, relates to the resistance of a material during the chewing process, with higher chewiness values indicating that the material is more difficult to chew. Table 1 summarizes these six parameters along with their units, description, and equations.

**Table 1:** Texture profile analysis. The texture profile analysis is based on double compression tests at *ε* = −50% peak strain, and uses the peak forces of the first and second loading cycles *F*_1_ and *F*_2_, the associated loading times *t*_1_ and *t*_2_, the areas under the loading paths *A*_1_ and *A*_3_, the areas under the unloading paths as *A*_3_ and *A*_4_, and the peak stress *σ*_1_ = *F*_1_/*A*, where *A* denotes the specimen cross section area. It extracts six parameters, the stiffness, hardness, cohesiveness, springiness, resilience, and chewiness that we summarize below with their units, descriptions, and equations.

| parameter | unit | description | equation |
| --- | --- | --- | --- |
| stiffness | kPa | slope of stress-strain curve during first compression when compressed to half of specimen height | $E = \frac{\sigma}{\varepsilon}$ |
| hardness | N | peak force at first compression when compressed to half of specimen height | $F_1$ |
| cohesiveness | – | material integrity during second loading/unloading cycle relative to first cycle, where 0 denotes complete disintegration | $\frac{A_3 + A_4}{A_1 + A_2}$ |
| springiness | – | material response time during second loading cycle relative to first cycle, recovery related to viscous properties | $\frac{t_2}{t_1}$ |
| resilience | – | material recovery from deformation during first unloading relative to first loading, 1 denotes elastic and $>1$ plastic behavior | $\frac{A_2}{A_1}$ |
| chewiness | N | material ease of biting<br>hardness times cohesiveness times springiness | $F_1 \frac{A_3 + A_4}{A_1 + A_2} \frac{t_2}{t_1}$ |

**Table 2:**
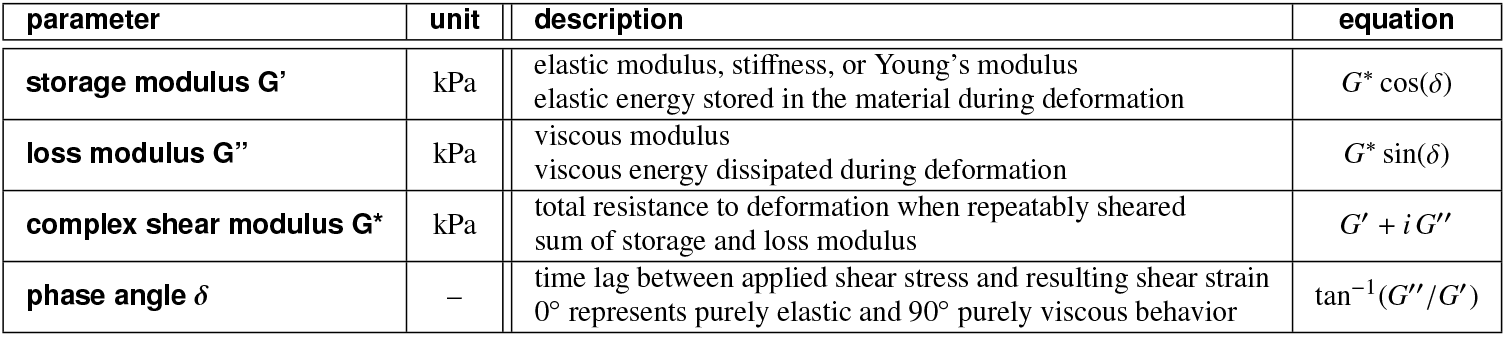
Rheological analysis. The rheological analysis is based on shear tests across a frequency sweep from 0.01-100 rad/s. It extracts four parameters, the storage modulus, loss modulus, complex shear modulus, and phase angle that we summarize below with their units, descriptions, and equations.

**Table 3:** Plant-based and animal meats. Products; brands; ingredients; texture profile analysis parameters stiffness, hardness, cohesiveness, springiness, resilience, and chewiness; and rheological parameters storage modulus, loss modulus, complex modulus, and tangent of phase angle. Results are reported as mean ± standard deviation from *n* = 13 samples for the texture profile analysis parameters and *n* = 5 samples for the rheology.

|  | AH animal hotdog | PH plant-based hotdog | AS animal sausage | PS plant-based sausage |
| --- | --- | --- | --- | --- |
| **brand** | Classic Uncured Wiener<br>Oscar Mayer, Kraft Heinz Co<br>Chicago, IL | Signature Stadium Hot Dog<br>Field Roast<br>Seattle, WA | Turkey Polska Kielbasa<br>Hallshire Farm<br>New London, WI | Vegan Frankfurter Sausage<br>Field Roast<br>Seattle, WA |
| **ingredients** | mechanically separated chicken, mechanically separated turkey, pork, water, corn syrup; contains 2% or less: salt, sodium phosphate, potassium chloride, sodium diacetate, sodium benzoate, sodium ascorbate, flavor, sodium nitrate | water, soybean oil, pea protein, potato starch, contains 2% or less: methylcellulose, carrageenan, sea salt, brown rice protein, faba bean protein, garlic powder, beet powder, cane sugar, natural flavor, wheat gluten, soy protein, konjac flour, potassium chloride, xanthan gum, spices, paprika, red rice flour | turkey, mechanically separated turkey, water, corn syrup; contains 2% or less: natural flavors, salt, dextrose, oat fiber, modified corn starch, monosodium glutamate, sodium erythorbate, yeast extract, sodium nitrite | filtered water, vital wheat gluten, expeller pressed safflower oil, organic expeller pressed palm fruit oil, barley malt, naturally flavored yeast extract, tomato paste, apple cider vinegar, paprika, sea salt, onions, spices, whole wheat flour |
| **stiffness** | 178.5 ± 20.58 kPa | 82.53 ± 13.79 kPa | 257.3 ± 24.08 kPa | 309.7 ± 32.58 kPa |
| **hardness** | 3.190 ± 0.332 N | 1.256 ± 0.182 N | 4.564 ± 0.675 N | 5.581 ± 0.717 N |
| **cohesiveness** | 0.804 ± 0.096 | 0.683 ± 0.178 | 0.621 ± 0.128 | 0.764 ± 0.076 |
| **springiness** | 0.809 ± 0.046 | 0.727 ± 0.081 | 0.747 ± 0.050 | 0.744 ± 0.053 |
| **resilience** | 1.114 ± 0.066 | 0.851 ± 0.078 | 1.067 ± 0.118 | 0.882 ± 0.048 |
| **chewiness** | 2.077 ± 0.360 N | 0.634 ± 0.244 N | 2.110 ± 0.510 N | 3.195 ± 0.707 N |
| **storage modulus** | 13.478 ± 0.934 kPa | 11.873 ± 0.923 kPa | 16.334 ± 1.225 kPa | 47.249 ± 4.187 kPa |
| **loss modulus** | 2.839 ± 0.156 kPa | 2.782 ± 0.305 kPa | 3.810 ± 0.283 kPa | 11.211 ± 1.456 kPa |
| **complex modulus** | 13.774 ± 0.946 kPa | 12.195 ± 0.968 kPa | 16.773 ± 1.287 kPa | 48.542 ± 4.410 kPa |
| **tangent of phase angle** | 0.210 ± 0.003 | 0.231 ± 0.008 | 0.233 ± 0.001 | 0.231 ± 0.010 |
|  | FT firm tofu | ET meats tofu | AT animal turkey | PT plant-based turkey |
| **brand** | Organic Firm Tofu<br>365 by Whole Foods Market<br>Austin, TX | Organic Tofu Extra Firm<br>House Foods<br>Garden Grove, CA | Spam Oven Roasted Turkey<br>Spam, Hormel Foods Co<br>Austin, MN | Ham-Style Roast<br>Tofurky<br>Hood River, OR |
| **ingredients** | water, organic soybeans, calcium sulfate, magnesium chloride | water, organic soybeans, calcium sulfate, calcium chloride | white turkey, turkey broth, salt, modified potato starch, sugar, dextrose, sodium nitrite | water, vital wheat gluten, tofu (water, soybeans, magnesium chloride, calcium chloride), expeller pressed canola oil; contains 2% or less: sea salt, spices, granulated garlic, cane sugar, natural flavors, natural smoke flavor, color, oat fiber |
| **stiffness** | 56.65 ± 14.06 kPa | 104.6 ± 14.17 kPa | 244.0 ± 16.70 kPa | 418.9 ± 41.72 kPa |
| **hardness** | 0.804 ± 0.334 N | 1.749 ± 0.333 N | 5.064 ± 0.649 N | 8.848 ± 0.821 N |
| **cohesiveness** | 0.783 ± 0.200 | 0.733 ± 0.096 | 0.657 ± 0.084 | 0.722 ± 0.080 |
| **springiness** | 0.752 ± 0.059 | 0.818 ± 0.063 | 0.805 ± 0.054 | 0.805 ± 0.055 |
| **resilience** | 0.891 ± 0.068 | 1.049 ± 0.141 | 0.920 ± 0.066 | 0.821 ± 0.058 |
| **chewiness** | 0.442 ± 0.131 N | 1.050 ± 0.257 N | 2.699 ± 0.630 N | 5.143 ± 0.829 N |
| **storage modulus** | 5.669 ± 0.499 kPa | 10.995 ± 1.017 kPa | 50.437 ± 4.131 kPa | 113.802 ± 10.383 kPa |
| **loss modulus** | 1.296 ± 0.077 kPa | 2.638 ± 0.175 kPa | 9.692 ± 0.704 kPa | 25.343 ± 2.998 kPa |
| **complex modulus** | 5.815 ± 0.503 kPa | 94.229 ± 0.858 kPa | 51.361 ± 4.189 kPa | 116.593 ± 10.785 kPa |
| **tangent of phase angle** | 0.229 ± 0.007 | 0.241 ± 0.007 | 0.192 ± 0.002 | 0.222 ± 0.006 |

**Figure 3:**
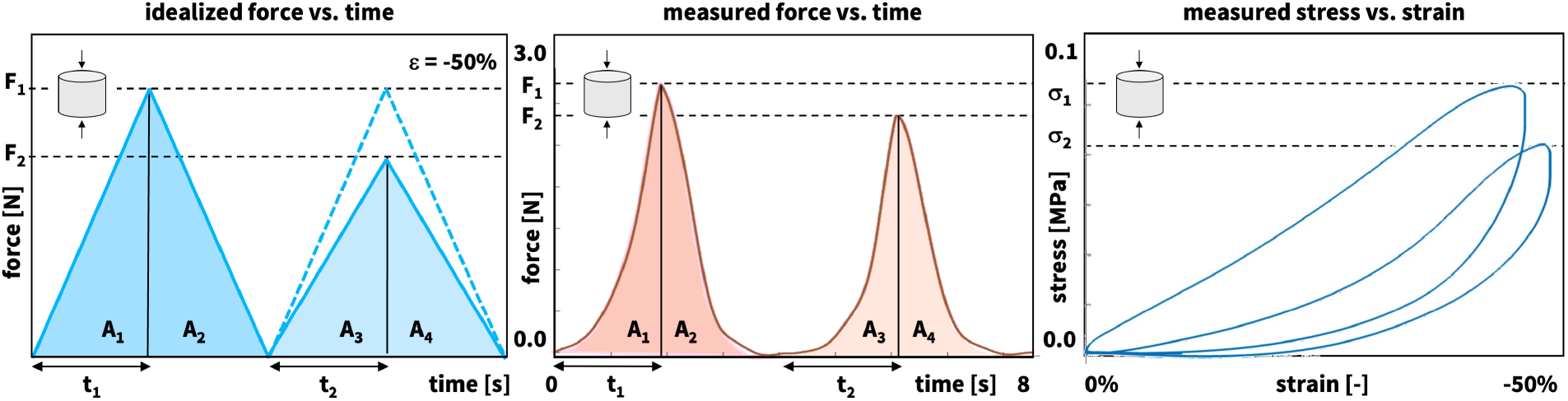
Texture profile analysis. For each of the eight meats, we perform *n* = 13 double compression tests with a peak compressive strain of *ε* = −50% and a strain rate of 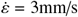. From the averaged force vs. time curves, we extract the peak forces of the first and second loading cycles *F*_1_ and *F*_2_, the associated loading times *t*_1_ and *t*_2_, the areas under the loading paths *A*_1_ and *A*_3_, and the areas under the unloading paths as *A*_3_ and *A*_4_. We convert the curves into stress vs. strain curves, where the stress *σ* = *F*/*A* is the recorded force *F* divided by the specimen cross section area *A*, and the strain 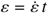 is the applied strain rate multiplied by the time *t*.

### 2.4. Rheological analysis

The shear tests serve as the basis for the rheological analysis. We conduct an *n* = 1 amplitude sweep to determine the strain for the frequency sweep. We perform *n* = 5 frequency sweep tests for each of the eight meats and average all five datasets from which we extract four rheological parameters. The *storage modulus G*′ = *G*^*^ cos(*δ*) is the elastic modulus, stiffness, or Young’s modulus that describes the elastic energy stored in the material during deformation. The *loss modulus G*′′ = *G*^*^ sin(*δ*) is the viscous modulus that describes the viscous energy dissipated during deformation. The *complex shear modulus G*^*^ = *G*′ + *i G*′′ is the sum of the storage and loss moduli and describes the total resistance to deformation as the material is repeatedly sheared. The *phase angle δ* = tan^−1^(*G*′′/*G*′) defines the time lag between applied shear stress and resulting shear strain and varies between 0^°^ ≤ *δ* ≤ 90^°^ where 0^°^ represents a purely elastic and 90^°^ a purely viscous behavior. Table 2 summarizes these four parameters along with their units, description, and equations.

### 2.5. Statistical analysis

We acquire all data using the TRIOS software and analyze the data sets in MATLAB. To quantify to which extent the plant-based meats differ from their animal counterparts, we perform a one-way analysis of variance ANOVA statistical test for each texture profile analysis or rheological parameter using MAT-LAB function anova1. If the ANOVA is significant, p < 0.05, we perform a Tukey-Kramer post-hoc test to determine which pairs of meats are significantly different from each other using the MATLAB function multcompare.

## 3. Results

We successfully perform double compression tests for the texture profile analysis and shear tests for the rheological analysis of five plant-based and three animal meats. Table 3 provides a comprehensive overview of the eight products, their brands and ingredients, their six textural parameters, and their four rheological parameters. Figures 4 and 6 summarize the pa-rameters of the texture profile analysis, the stiffness, hardness, cohesiveness, springiness, resilience, and chewiness, and the parameters of the rheological analysis, the storage, loss, and complex shear modulus, and the phase angle, all in terms of box and whisker plots that highlight the minimum, lower quartile, median, upper quartile, maximum values, and outliers for all eight meats. Table 4 summarizes the textural difference between the plant-based and animal meats for hotdog, sausage, turkey, and, for comparison, for firm and extrafirm tofu. Figures 5 and 7 summarize the textural and rheological difference across all eight meats.

**Table 4:** Textural difference between plant-based and animal meats. Results of one-way ANOVA tests with Tukey-Kramer correction for similar meats, animal vs plant-based hotdog, animal vs plant-based sausage, and animal vs plant-based turkey, and, for comparison, firm and meats tofu. Stars denote statistically significant differences in textural properties, where * denotes p-value < 0.05, ** denotes p-value < 0.01, *** denotes p-value < 0.001, and **** denotes p-values < 0.0001.

|  | AH vs PH<br>plant based vs animal<br>hotdog | AS vs PS<br>plant based vs animal<br>sausage | AT vs PT<br>plant based vs animal<br>turkey | FT vs ET<br>firm vs meats<br>tofu |
| --- | --- | --- | --- | --- |
| stiffness | **** | **** | **** | **** |
| hardness | **** | **** | **** | **** |
| cohesiveness |  |  |  |  |
| springiness | * |  |  |  |
| resilience | **** | **** |  | *** |
| chewiness | **** | **** | **** |  |

**Figure 4:**
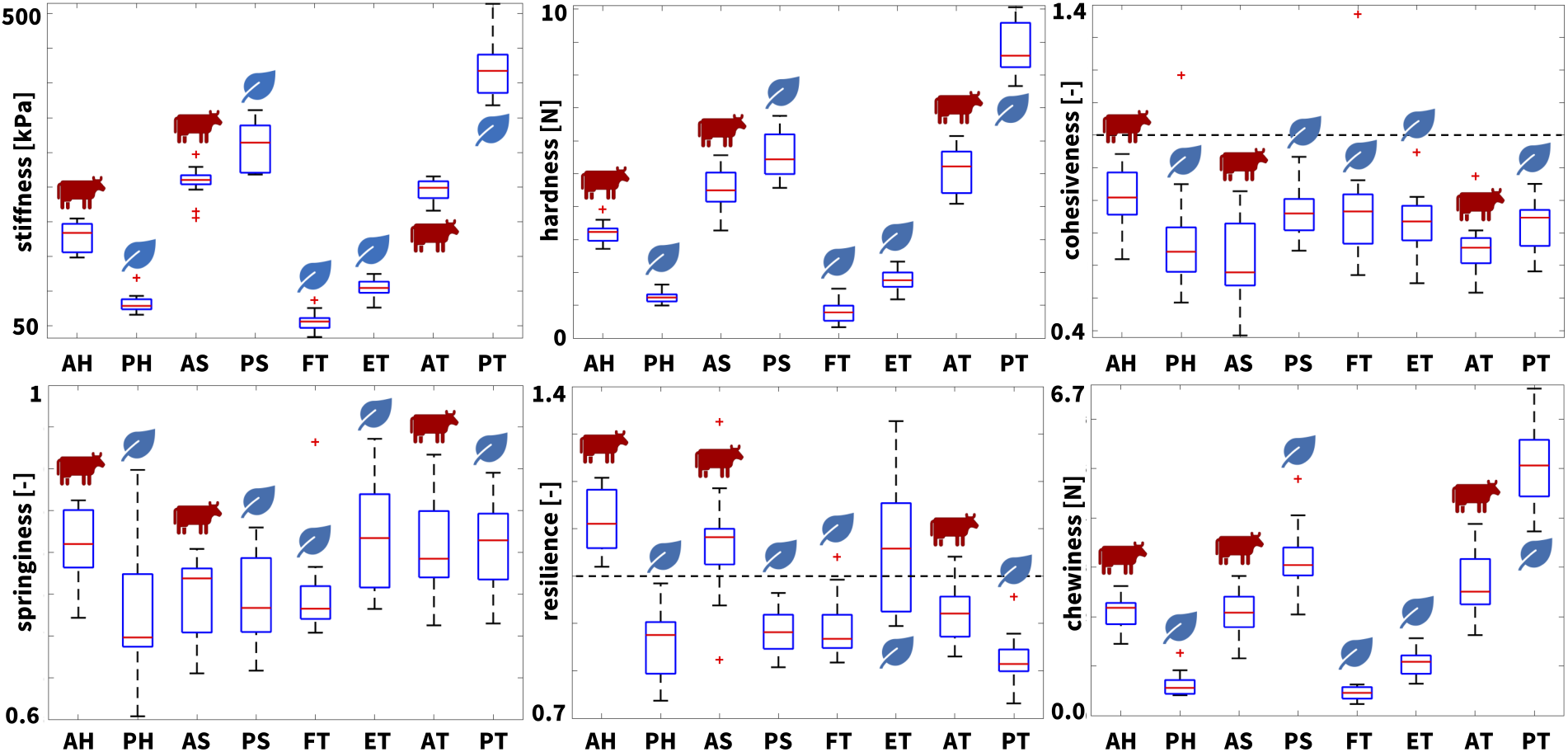
Texture profile analysis. Stiffness, hardness, cohesiveness, springiness, resilience, and chewiness of all eight meats; the box and whisker plots summarize the minimum, lower quartile, median, upper quartile, maximum values, and outliers across *n* = 13 double compression tests; AH animal hotdog, PH plant-based hotdog, AS animal sausage, PS plant-based sausage, FT firm tofu, ET meats tofu, AT animal turkey, PT plant-based turkey.

**Figure 5:**
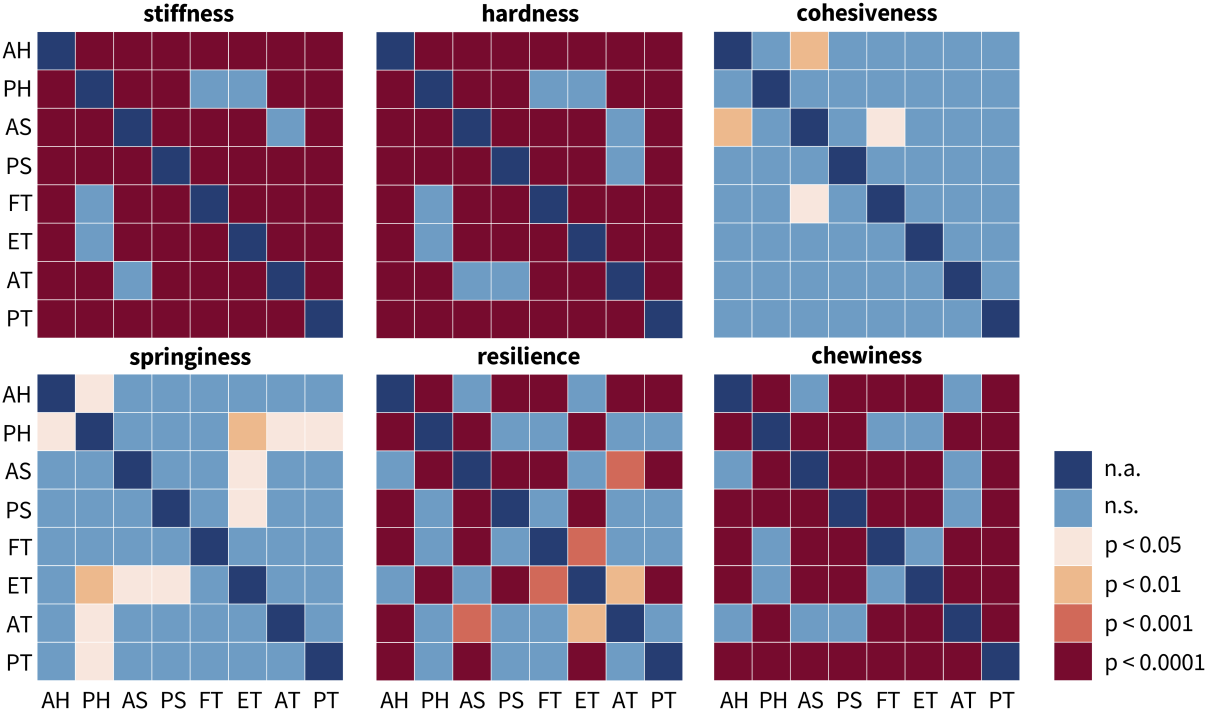
Textural difference across all eight meats. Results of one-way ANOVA tests with Tukey-Kramer correction for texture profile analysis of all eight meats in terms of stiffness, hardness, cohesiveness, springiness, resilience, and chewiness. Color code denotes statistically significant differences in texture analysis profiling, where blue denotes no significance, light orange denotes p-values < 0.05, orange denotes p-values < 0.01, red denotes p-values < 0.001, and dark red denotes p-values < 0.0001; AH animal hotdog, PH plant-based hotdog, AS animal sausage, PS plant-based sausage, FT firm tofu, ET meats tofu, AT animal turkey, PT plant-based turkey.

**Figure 6:**
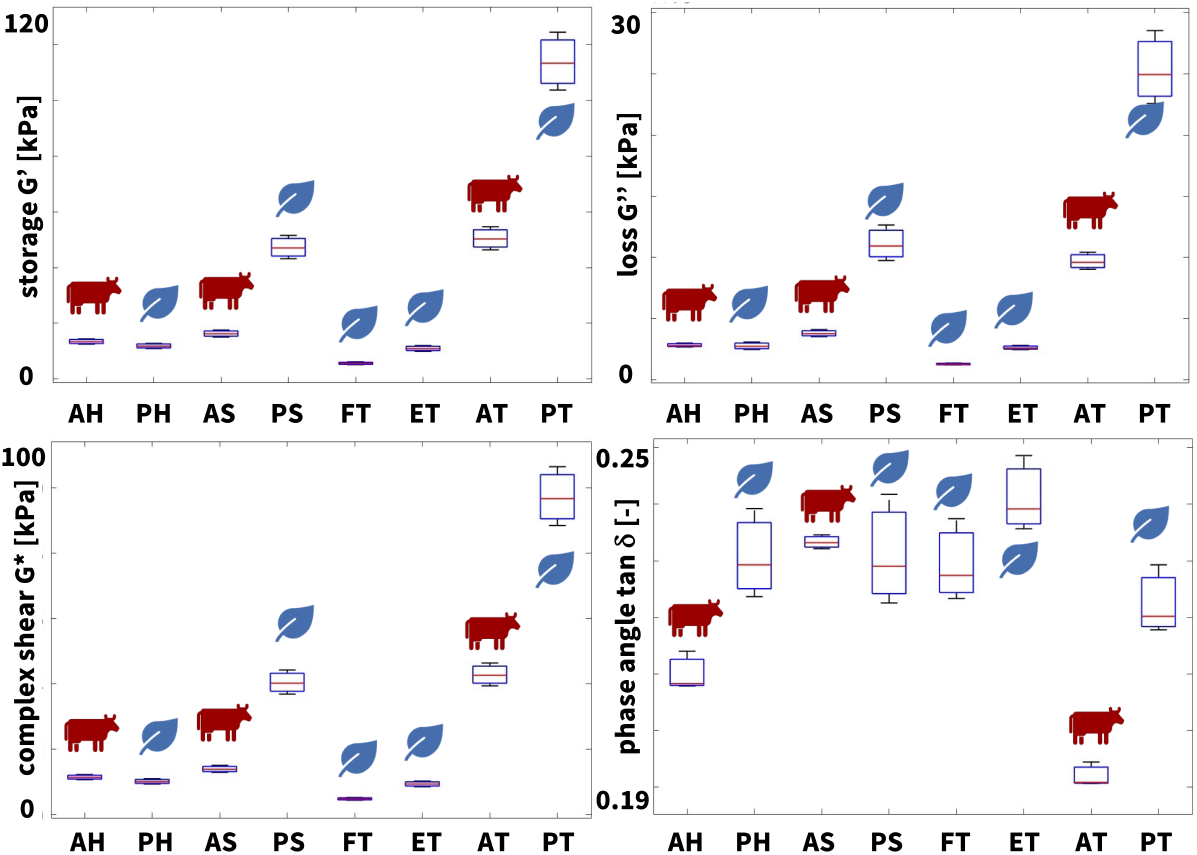
Rheological characterization. Storage modulus, loss modulus, complex shear modulus, and phase angle of all eight meats; the box and whisker plots summarize the minimum, lower quartile, median, upper quartile, maximum values, and outliers across *n* = 5 frequency sweep shear tests; AH animal hotdog, PH plant-based hotdog, AS animal sausage, PS plant-based sausage, FT firm tofu, ET meats tofu, AT animal turkey, PT plant-based turkey.

**Figure 7:**
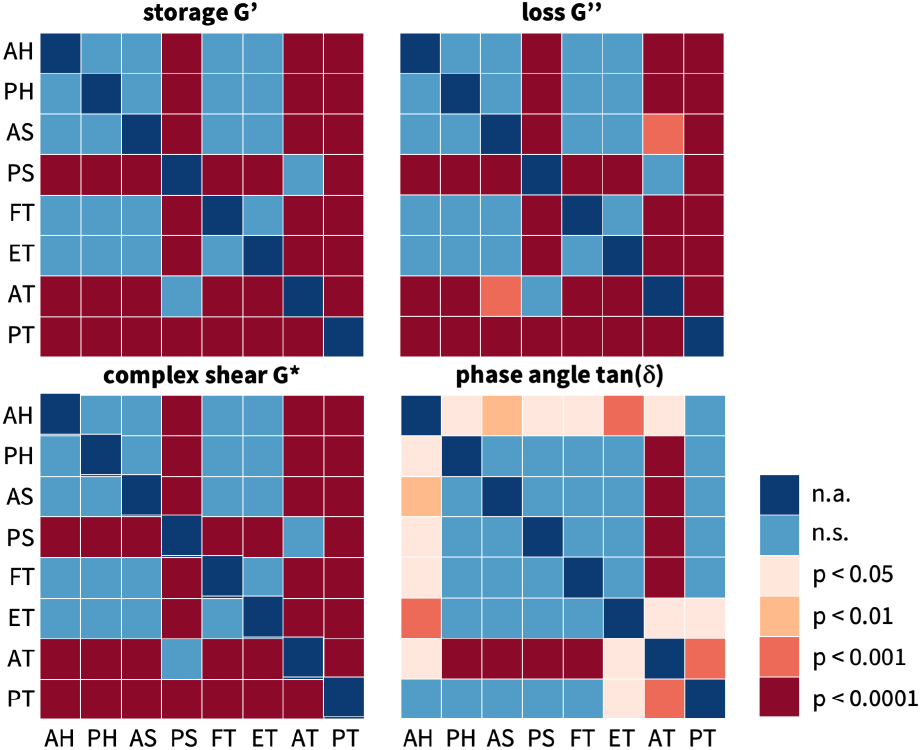
Rheological differences across all eight meats. Results of one-way ANOVA tests with Tukey-Kramer correction for rheological testing of all eight meats in terms of storage modulus, loss modulus, complex shear modulus, and tangent. Color code denotes statistically significant differences in texture analysis profiling, where blue denotes no significance, light orange denotes p-values < 0.05, orange denotes p-values < 0.01, red denotes p-values < 0.001, and dark red denotes p-values < 0.0001; AH animal hotdog, PH plant-based hotdog, AS animal sausage, PS plant-based sausage, FT firm tofu, ET meats tofu, AT animal turkey, PT plant-based turkey.

### Our recordings agree well with reported results

Our recordings for animal sausage and animal turkey agree well with a previous study that reported stiffnesses on the order of 90-120 kPa, hardnesses on the order of 5-8 kPa, cohesivenesses on the order of 0.78-0.85, springinesses on the order of 0.83-0.87, chewinesses on the order of 3.7-4.8, and re-siliences on the order of 1.4-1.6 and storage moduli *G*′ of 25-30 kPa, loss moduli *G*′′ of 7 kPa, complex shear moduli *G*^*^ of 25-30 kPa, and phase angles tan *δ* of 0.18.5 to 0.192 [24]. Surprisingly, although our tested sausage and turkey products are not identical to these products, our values all fall well within these ranges. We have previously tested the identical products as in this study, but only for their elastic properties [31]. Strikingly, we found a same order in the compression stiffness, with plant-based turkey, plant-based sausage, animal turkey, and animal sausage testing the stiffest, and animal hotdog, meats tofu, plant-based hotdog, and firm tofu testing the softest [31]. Notably, these previous stiffnesses, averaged over tension, compression, and shear tests, lie square within the ranges of the double compression stiffnesses and frequency sweep storage moduli of the present study. Yet, in addition, the present study reveals some exciting new insights.

### Plant-based meats successfully replicate the elastic properties of processed animal meat

From the loading branch of the first load cycle of the double compression test, we can extract the elastic *stiffness* as the ratio of peak stress to peak strain, here at 50% compression. Since the stress-strain relations are non-linear, we approximate the stiffness across the entire loading interval using least-squares regression. Figure 4, top left, illustrates the stiffnesses for all eight meats and Table 3 summarizes the stiffness values with means and standard deviations. Interestingly, the turkey and sausage products consistently display stiffnesses larger than 200 kPa, including plant-based turkey with 418.9 kPa, plant-based sausage with 309.7 kPa, animal turkey with 244.0 kPa, and animal sausage with 257.3 kPa. The hotdog and tofu products display stiffnesses smaller than 200 kPa, including animal hotdog with 178.5 kPa, meats tofu with 104.6 kPa, plant-based hotdog with 82.53 kPa, and firm tofu with 56.65 kPa. Against these trends, the stiffnesses of the plant-based and animal products of the same categories are statistically significantly different, including animal vs. plant-based hotdog (p < 0.0001), animal vs. plant-based sausage (p < 0.0001), and animal vs. plant-based tofu (p < 0.0001), see Figure 5 and Table 4.

We can also use the loading branch of the first load cycle of the double compression test to extract the *hardness* as the peak force, in our case at 50% compression. Figure 4, top middle, illustrates the hardnesses for all eight meats and Table 3 summarizes the hardness values with means and standard deviations. As expected, the four turkey and sausage products that display the largest stiffnesses have the largest hardnesses, all larger than 4 N, including plant-based turkey with 8.8 N, plant-based sausage with 5.6 N, animal turkey with 5.1 N, and animal sausage with 4.6 N. Similarly, the four hotdog and tofu products that display the smallest stiffnesses have hardnesses smaller than 4 N, including animal hotdog with 3.2 N, plant-based hot-dog with 1.3 N, meats tofu with 1.7 N, and firm tofu with 0.8 N. Similar to the stiffnesses, the hardnesses of the plant-based and animal products of the same categories are statistically significantly different, including animal vs. plant-based hotdog (p < 0.0001), animal vs. plant-based sausage (p < 0.0001), and animal vs. plant-based tofu (p < 0.0001), see Figure 5 and Table 4. Notably, for a perfectly linear elastic behavior, we can estimate the hardness from the stiffness values. Our samples have a diameter of 8 mm and a cross sectional area 50.3 mm^2^.

This implies that, for ideal linear elasticity, with a strain of 0.5, a stiffness value in kilopascals translates into 40 times a hardness value in Newtons. For our measured stiffnesses and hardnesses, this factor is indeed close to 40: It varies from 48.2 for animal turkey and 47.3 for plant turkey, suggesting that these products are almost perfectly linearly elastic, to 65.7 for plant hotdog and 70.5 for firm tofu, indicating that these products are non-linearly elastic.

To confirm our observations associated with the elastic stiffness of the texture profile analysis, we can extract the *storage modulus* from the frequency sweep shear test. Figure 6, top left, illustrates the storage moduli for all eight meats and Table 3 summarizes the storage moduli with means and standard deviations. Our rheological measurements confirm the texture profile analysis, with the turkey and sausage products consistently displaying the largest storage moduli, including plant-based turkey with 113.8 kPa, animal turkey with 50.4 kPa, plant-based sausage with 47.2 kPa, and animal sausage with 16.3 kPa. Sim-ilarly, the hotdog and tofu products display the smallest storage moduli, including animal hotdog with 13.5 kPa, plant-based hotdog with 11.9 kPa, meats tofu with 11.0 kPa, and firm tofu with 5.7 kPa. Theoretically, for isotropic materials, the storage modulus *G*′ relates to the Young’s modulus or elastic stiffness as *G*′ = 2 (1 + ν) *E*, where ν is Poisson’s ratio. If we assume that our meats are incompressible, with ν = 0.5, the storage modulus would be three times the stiffness, *G*′ = 3 *E*. While our storage moduli are generally smaller than three times the stiffness, we note that this discrepancy is consistent across all meats, and likely associated with the different loading rates. At the same time, we note that the stiffness ranking is the same for both measurements: The meats with the largest stiffnesses, the turkey and sausage products, also display the largest storage moduli.

In summary, the elastic properties of plant-based and animal meat vary by an order of magnitude ranging from 418.9 kPa for the plant-based product Ham-Style Roast Tofurky to 56.65 kPa for Organic Firm Tofu, while the three animal products, Spam Oven Roasted Turkey with 244.0 kPa, Turkey Polska Kielbasa with 257.3 kPa, and Classic Uncured Wiener with 178.5 kPa, rank in between these two extremes. In food science, ingredients such as vital wheat gluten, soy protein isolate, collagen, transglutaminase, egg whites, and hydrocolloids such as carrageenan, xanthan gum are commonly used to increase the elasticity of processed meat products. Table 3 confirms that these ingredients are present in the three plant-based meats: Plant-based hotdog contains wheat gluten, soy protein, and xanthan gum, and plant-based sausage and plant-based turkey contain vital wheat gluten. These ingredients enhance the elasticity by creating robust, flexible protein networks that give the meat product a firmer, stretchier texture. *Taken together, these observations suggest that modern food fabrication techniques can comfortably create plant-based meats that replicate the full range of elastic textures of processed animal meat*.

### Plant-based meats replicate the loss of integrity of processed animal meat ranging from 18% to 38%

Beyond the elastic behavior, our texture profile analysis provides valuable insight into the inelastic behavior of plant-based and animal meat. From the comparison of the first and second cycles of the double compression test, we can extract the *cohesiveness* as the ratio of the area under the second load cycle compared to the first, (*A*_3_ + *A*_4_)/(*A*_1_ + *A*_2_). Figure 4, top right, illustrates the cohesiveness for all eight meats and Table 3 summarizes the cohesiveness with means and standard deviations. The cohesiveness is a measure of dissipation. It characterizes the loss of material integrity, where a value of one indicates a completely intact material associated with a purely elastic response and a value of zero indicates complete material disintegration. As expected, all eight meats display a mean cohesiveness smaller than one, i.e., *A*_1_ + *A*_2_ > *A*_3_ + *A*_4_, ranging from 0.804 for animal hotdog that stayed most intact to 0.621 for animal sausage that disintegrated the most. Notably, all plant-based products, with 0.764 for plant-based sausage, 0.722 for plant-based turkey, and 0.683 for plant-based hotdog, rank somewhere in between these two extremes.

Another parameter that we can extract from the comparison of the first and second loading cycles is the *springiness*, the ratio of the second loading time compared to the first, *t*_2_/*t*_1_, which is equivalent to the ratio of the second compression distance to the first, *d*_2_/*d*_1_, given a constant strain rate. Importantly, this is the time that the rheometer is in contact with the sample, not simply the duration of compressive loading. As a result, a springiness value of one indicates a perfectly elastic response for which the sample returns to its original height. Figure 4, bottom left, illustrates the springiness for all eight meats and Table 3 summarizes the springiness with means and standard deviations. Across all eight meats, the springiness is consistently less than one, *t*_2_ < *t*_1_, ranging from 0.727 for plant-based hotdog to 0.818 for extrafirm tofu. Notably, unlike the elastic properties, the cohesiveness and springiness of all meats fall into a rather narrow range and very few combinations are statistically significantly different, see Figure 5. From Table 4, only the springiness of animal hotdog is significantly different from plant hotdog with p<0.5.

From the comparison of the first loading and unloading curve of the double compression test, we can extract the *resilience* as the ratio of the area under unloading curve compared to the loading curve, *A*_2_/*A*_1_. Figure 4, bottom middle, illustrates the resilience for all eight meats and Table 3 summarizes the resilience with means and standard deviations. The resilience is a measure of material recovery, where a value of one indicates a purely elastic response and a value less than one indicates plastic damage. Unexpectedly, animal hotdog, animal sausage, and extrafirm tofu display a mean resilience larger than one. In the raw data, these meats exhibited a non-linear unloading curve with a bump or plateau as we can see in Figure 3, resulting in *A*_2_ > *A*_1_. As expected, for all plant-based products with the exception of extrafirm tofu, the resilience is less than one, i.e., the unloading area is less than the loading area, *A*2 < *A*1. With plant-based turkey having the lowest resilience at 0.821, this suggests that all meats are able to withstand 50% compression without completely disintegrating, which is further evidenced by the cohesiveness values of these meats all above 0.6.

From the product of the hardness, cohesiveness, and springiness, we can extract another texture parameter, the *chewiness* that characterizes the ease of biting. Since the springiness and cohesiveness parameters across all meats are similar to each other with very few combinations significantly different, the chewiness is dominated by the hardness, the peak force of the first loading cycle. Accordingly, the least difficult products to chew are plant-based hotdog and both firm and extrafirm tofu, products with the smallest hardness values, with plant-based hotdog with a chewiness of 0.6 N, firm tofu with 0.4 N, and extrafirm tofu with 1.1 N. The hardest products to chew are plant-based sausage with 3.2 N and plant-based turkey with 5.1 N. Interestingly, all animal meats fall within the chewiest and least chewy plant-based products, with animal hotdog with 2.1 N, animal sausage with 2.1 N, and animal turkey with 2.7 N. *Taken together, probably the most notable observation associated with the inelastic parameters of springiness and cohesiveness is that all eight meats consistently lose between 18% to 38% of integrity during the first loading cycle of 50% compression*. All plant-based products are engineered to fall well within this range–no product remains entirely elastic and no product degrades entirely.

### Plant-based meats successfully replicate the viscous properties of processed animal meat

Our rheological analysis provides additional insight into the viscous behavior of plant-based and animal meat. From the frequency sweep shear test, we can extract the *loss modulus* as a parameter to characterize the viscous behavior. Figure 6, top right, illustrates the loss moduli for all eight meats and Table 3 summarizes the loss moduli with means and standard deviations. Interestingly, similar to the storage moduli *G*′, the turkey and sausage products also display the largest loss moduli *G*′′, including plant-based turkey with 25.3 kPa, plant-based sausage with 11.2 kPa, animal turkey with 9.7 kPa, and animal sausage with 3.8 kPa. Similarly, the hotdog and tofu products display the smallest loss moduli, including animal hotdog with 2.8 kPa, plant-based hotdog with 2.8 kPa, extrafirm tofu with 2.6 kPa, and firm tofu with 1.3 kPa. Remarkably, with the exception of plant-based sausage and animal turkey, the order of the loss or viscous moduli is identical to the order of the storage or elastic moduli.

This order naturally translates into the *complex shear modulus*, the sum of the storage and loss moduli, *G*^*^ = *G*′ + *i G*′′, and the the *complex shear moduli* for all eight meats in Figure 6, bottom left, look fairly similar to the storage and loss moduli in Figure 6, top left and right. The complex shear moduli range from 116.6 kPa for plant-based turkey to 5.8 kPa for firm tofu, with the animal products in between these two extremes.

The last rheological parameter is the *phase angle δ*, which we can extract as the ratio of the loss and storage moduli, tan *δ* = *G*′′/*G*′. Figure 6, bottom right, illustrates the phase angles for all eight meats and Table 3 summarizes the phase angles with means and standard deviations. Unlike the storage, loss, and complex shear moduli, the phase angle lies in a very narrow range, varying only from 0.192 or 10.9^°^ for extrafirm tofu to 0.241 or 13.5^°^ for animal turkey. For an ideally elastic behavior, tan *δ* = 0, *δ* = 0^°^, and *G*′′ = 0 kPa, while for an ideally viscous behavior, tan *δ* = ∞, *δ* = 90^°^, and *G*′ = 0 kPa. This suggests that the behavior of all eight meats is more elastic than viscous.

Food scientists manipulate the viscosity of processed meats by adding ingredients like starches such as potato starch or modified corn starch, hydrocolloids such as carrageenan or xanthan gum, proteins such as soy protein or gluten, and binders such as methylcellulose [5]. Table 3 confirms that these ingredients are indeed present in the three plant-based meats: plant-based hotdog contains potato starch, xanthan gum, and methyl-cellulose, and plant-based sausage and plant-based turkey contain vital wheat gluten. Notably, starches are even present in two of the animal products: animal sausage contains corn starch and animal turkey contains potato starch. These ingredients increase the viscosity by absorbing water, forming gels, or creating cohesive matrices that enhance the thickness and stability of the meat mixture. *Taken together, our findings suggest that modern food fabrication techniques can reliably create comminuted plant-based meats that closely mimic the narrow range of the viscous properties of processed animal meat*.

### Texture profile analysis and rheology are predictors of our perception of taste

Leading up to this study, we conducted a survey of *n* = 16 participants who ate all eight meats and rated them by twelve sensory attributes including soft, hard, brittle, chewy, gummy, viscous, springy, sticky, fibrous, fatty, moist, and meaty [21, 35], on a scale from one to five [31]. The study was reviewed and approved by the Institutional Review Board at Stanford University under the protocol IRB-75418. Figure 8 shows that the participants ranked animal and plant hotdog and animal and plant sausage all similarly to each other for softness, hardness, and chewiness. In contrast, they ranked the plant-based products as less fatty, less moist, and less meaty than their animal versions. Plant turkey was more hard and chewy, but less fatty, moist, and meaty than animal turkey. The firm and extrafirm tofu products were the softest, least fatty, and least meaty of all eight products [31]. Most notably, the hardnesses from our text profile analysis in Figure 4 generally reflect the rankings from the food texture survey in Figure 8, with plant turkey being the hardest and both tofus being the softest. The stiffnesses from the text profile analysis in Figure 4 and the storage moduli from the rheology in Figure 6 confirm these trends: They are closely correlated to the sensory perception of hard and inversely correlated to the perception of soft in Figure 8. Interestingly, plant based turkey was also perceived the most chewy in Figure 8, in agreement with the texture profile analysis in Figure 4. *Taken together, our findings suggest that both texture profile analysis and rheology generally align with our sensory perception of texture but provide a more objective and more reproducible characterization than food texture surveys*.

**Figure 8:**
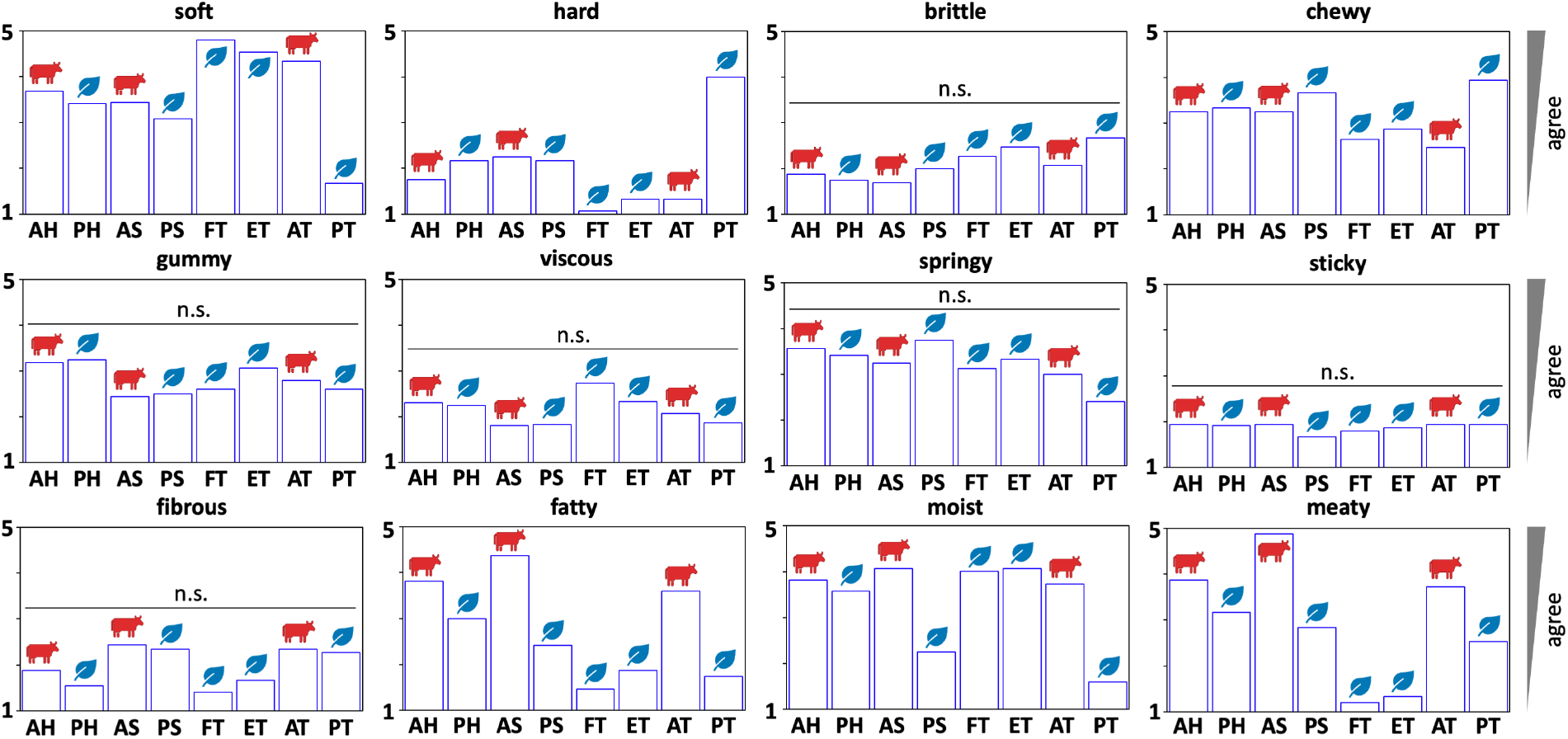
Food texture survey. Soft, hard, brittle, chewy, gummy, viscous, springy, sticky, fibrous, fatty, moist, and meaty scores of all of all eight meats; the box plots summarize the mean scores of the texture survey across *n* = 16 participants [31]; all participants ate cooked samples of all eight products, and ranked their texture features on a 5-point Likert scale ranging from 5 for strongly agree to 1 for strongly disagree; AH animal hotdog, PH plant-based hotdog, AS animal sausage, PS plant-based sausage, FT firm tofu, ET meats tofu, AT animal turkey, PT plant-based turkey.

### Critical comparison of texture profile analysis and rheology

Texture profile analysis is a double compression test from which we can derive parameters such as hardness, springiness, and chewiness that we can correlate to sensory tests [14, 23, 31]. While this test is simple to perform and easy to analyze, it suffers from a lack of standardization across research groups, inconsistent definitions of how to calculate the output parameters, and general confusion over mutually exclusive parameters [32]. Without a set standard, individual researchers must choose their own amount of compression, rate of compression, and sample dimensions, which strongly limits cross-study comparisons. As a result, stiffness is the most meaningful parameter to report [25]. Its definition is scaled by the specimen dimensions, and it is a strong indicator of elasticity provided the deformation is nearly linear. In contrast, hardness, the maximum recorded force of the first compression, depends on the amount of compression, the rate of compression, and the cross section area, so it provides no additional information than stiffness.

Cohesiveness is a measure of inelasticity or dissipation, and it characterizes the loss of integrity of the sample. A perfectly elastic material would have a cohesiveness of one, any plastic deformation or disintegration would result in a value less than one. However, in our data, we observe samples where the second compression peak is recorded as higher than the first peak, which may be an effect from the rheometer sampling rate, sample slippage, buckling, or moisture content change. For a perfectly elastic material, springiness would always equal one given a constant strain rate and peak compressive strain. For a plastically deformed material, after the first compression, the springiness value would be less than one. Resilience is another measure of inelasticity or dissipation, somewhat similar to cohesiveness, and should also be less than or equal to one. Again, our data display several instances where the area associated with decompression is larger than the area associated with compression. In most of these cases, we observe a second mini-peak during the decompression phase, which could either be an artifact of the rheometer failing to record at high rates or the sample changing shape or slipping. Chewiness is a derived property as the product of hardness, cohesiveness, and springiness; if the latter two are close to one, then chewiness is closely related to hardness and not an independent parameter. Taken together, our data showcase the limitations of performing a texture profile analysis with a standard rheometer on materials with non-linear, complex behaviors.

In contrast to texture profile analysis, the parameters from rheology are well-defined metrics with a clear mechanical interpretation. The storage modulus characterizes the elastic behavior and the loss modulus characterizes the viscous behavior. We find that the storage modulus reflects the stiffness of the meats, with firm and extrafirm tofu having the lowest stiffnesses and lowest storage moduli, and with plant turkey having the highest storage modulus and highest stiffness. *Taken together, the stiffness of the first compression test and the storage and loss moduli of the rheology are the most meaningful and reliable parameters to report when characterizing the mechanics of meat with a standard rheometer*.

## Conclusion

Plant-based meats have a lower environmental impact than animal meats. Convincing people to switch to these products remains a challenge, in part because plant-based meats have a different texture and mouth-feel than animal meat. Here we characterize the mechanical differences between five plant-based and three comparable animal meats using texture profile analysis form double compression tests and rheology from frequency sweeps. We observed inconsistencies when interpreting the parameters from the texture profile analysis, especially for non-linear complex data. In contrast, the stiffness modulus from the first compression and the storage and loss moduli from rheology, three well-understood and widely-used metrics, allow us to directly compare the mechanical behavior of plant-based and animal meats and correlate it to our sensory perception. From these comparisons, we conclude that the stiffest meats, plant-based and animal turkey and sausage, consistently have the highest storage and loss moduli. Plant-based turkey with stiffness and storage moduli of 419 kPa and 114 kPa is the stiffest product, and tofu with 57 kPa and 6 kPa is the softest; while all animal meat products fall within these two extremes. These observations agree with our sensory perception where participants of a food texture survey ranked the plant-based turkey product the hardest and least soft and the tofu products the softest and least hard. These findings suggest that both texture profile analysis and rheology are valuable quantitative and objective predictors of our qualitative and subjective perception of texture. More broadly, our study suggests that, with the right ingredients, additives, and formulation, modern food fabrication techniques can comfortably create plant-based meats that replicate the full spectrum of elastic textures and the narrow range of viscous properties of processed animal meat.

## Data Availability

Raw data will be available upon request. Table 3 includes all derived parameters.

## Acknowledgments

This work was supported by an NSF Graduate Student Fellowship to Reese Dunne, Ethan Darwin, and Skyler St. Pierre, a Stanford Graduate Fellowship to Reese Dunne, a GEM Fellowship to Valerie Perez Medina, a Stanford DARE Fellowship to Skyler St. Pierre, seed funding from the Stanford Plant-Based Diet Initiative and from Food System Innovations, the NSF CMMI Award 2320933 *Automated Model Discovery for Soft Matter* and the ERC Advanced Grant 101141626 *DISCOVER* to Ellen Kuhl.

## Notes

### Competing Interest Statement

The authors have declared no competing interest.

### Summary of Updates

slight edits of typos, addition of means and standard deviations to the data table

